# HLA polymorphism impacts immune response to neoepitopes and survival in APOBEC-mutated cancers

**DOI:** 10.1101/2024.08.07.607038

**Authors:** Faezeh Borzooee, Alireza Heravi-Moussavi, Mani Larijani

## Abstract

APOBEC3A and APOBEC3B genome mutators drive tumor evolution and drug resistance but may also generate neoepitopes for cytotoxic T cells (CTL). Given the extensive polymorphism of Class I HLA, the CTL immunopeptidome, comprised of all 8-11mer peptides presented by an individual’s six HLA class I alleles, varies person-to-person. We predicted the genome-wide impact of APOBEC3A/B-driven mutations on the immunogenicity of the immunopeptidomes of several thousand class I HLA alleles. Analysis of several billion APOBEC3-mediated mutations revealed that HLA class I alleles vary markedly in the susceptibility of their immunopeptidome to mutations. A subset of alleles of A1-A3 and B44 supertype supported increased neoepitopes. Notably, the immunogenicity changes supported by an individual’s HLA class I alleles in response to APOBEC3 mutations predict survival in APOBEC3-mutated tumors and correlate with CTL activation. Thus, immunogenicity changes mediated by APOBEC3s impact survival, making HLA class I genotype a prognostic marker in APOBEC3-mutated tumors.

## Background

The APOBEC3 cytidine deaminases are a major source of tumor genome mutations. APOBEC3s can mutate any cytidine, but each member has a favored hotspot. APOBEC3A (A3A) and APOBEC3B (A3B) which respectively prefer 5’YT<u>C</u>A3’, 5’RT<u>C</u>A3’ (R=purine, Y=pyrimidine) have emerged as the major mutators responsible for single base substitution (SBS) 2 and 13 signatures in solid tumors including lung, breast, bladder, head and neck and ovarian cancer ^1–6^. A3A- and A3B promote tumorigenesis, tumor clonal heterogeneity and drug resistance in cell culture and mouse models, and correlate with poor prognosis in human cancers ^6–14^

Central to anti-tumor immunity are CD8^+^ cytotoxic T cells (CTL) whose T cell receptor (TCR) recognizes 8-11mer peptide epitopes presented on tumour cells by MHC (encoded by HLA) class I. Established as A3A and A3B are as pro-tumorigenic factors, several findings suggest that they may also promote anti-tumor immunity: first, A3B-mediated generation of heteroclitic or neo- epitopes for CTL has been demonstrated in a mouse model ^15^. Second, higher expression of A3B and A3G in cervical cancer correlates with higher immune cell tumor infiltration ^16,17^. Third, a bioinformatics predictive study suggests that APOBEC3-mediated SBS2/13 mutations increase the hydrophobicity of CTL epitopes which promotes TCR and HLA binding ^18,19^.

HLA class I are highly polymorphic comprised of several thousand alleles. The HLA polymorphisms particularly affect their peptide epitope binding “anchor” residues, culminating in HLA alleles presenting different sets of peptides. Thus, each person’s haplotype of six A, B, C alleles determine a unique CTL immunopeptidome. Accordingly, HLA allelic differences have been associated with anti-tumour immunity and immunotherapy outcomes ^20,21^.

In contrast to promotion of CTL epitope generation, APOBEC3-mediated mutations may also diminish peptide epitope immunogenicity leading to tumor CTL evasion. To gain insight into the balance of these opposing outcomes, we examined the predicted impact of A3A- or A3B-mediated mutations on HLA class I anchor and TCR contact amino acid positions for the CTL immunopeptidomes of several thousand HLA I alleles.

We report that A3A- and A3B-mediated genome mutations can indeed enhance or diminish CTL epitope immunogenicity. The relative probability of these two outcomes varies infinitely across the immunopeptidomes restricted to different HLA class I alleles, and the carriage of HLA alleles whose immunopeptidomes predominantly gain or lose immunogenicity due to A3A- and A3B- mediated mutations predicts survival in APOBEC3-mutated cancers. These results underscore the intricate relationship between APOBEC3-mediated mutations, HLA genotype and anti-tumor immunity, and highlight the importance of considering individual immunogenetic variations in APOBEC3-mutated tumors.

## Results

### Genome-wide prediction of APOBEC3-mediated mutation impact on the HLA Class I immunopeptidome

Figure 1 illustrates the overall workflow of this study. We simulated all possible mutations in hotspot motifs of A3A (YT<u>C</u>) and A3B (RT<u>C</u>) on both strands of the genomic sequences encoding each peptide in the CTL immunopeptidome, which includes all possible 8-11mer peptides that can result from proteasomal degradation of the human proteome and potentially be presented by at least one of 2910 HLA class I. In contrast to A3A and A3B which are the main mutational culprits in solid tumors, AID’s mutagenic activity is largely restricted to leukemia/lymphomas for which the CTL response is not a key factor. Thus, mutations in AID hotspots (WRC) were simulated as a control.

**Figure 1.**
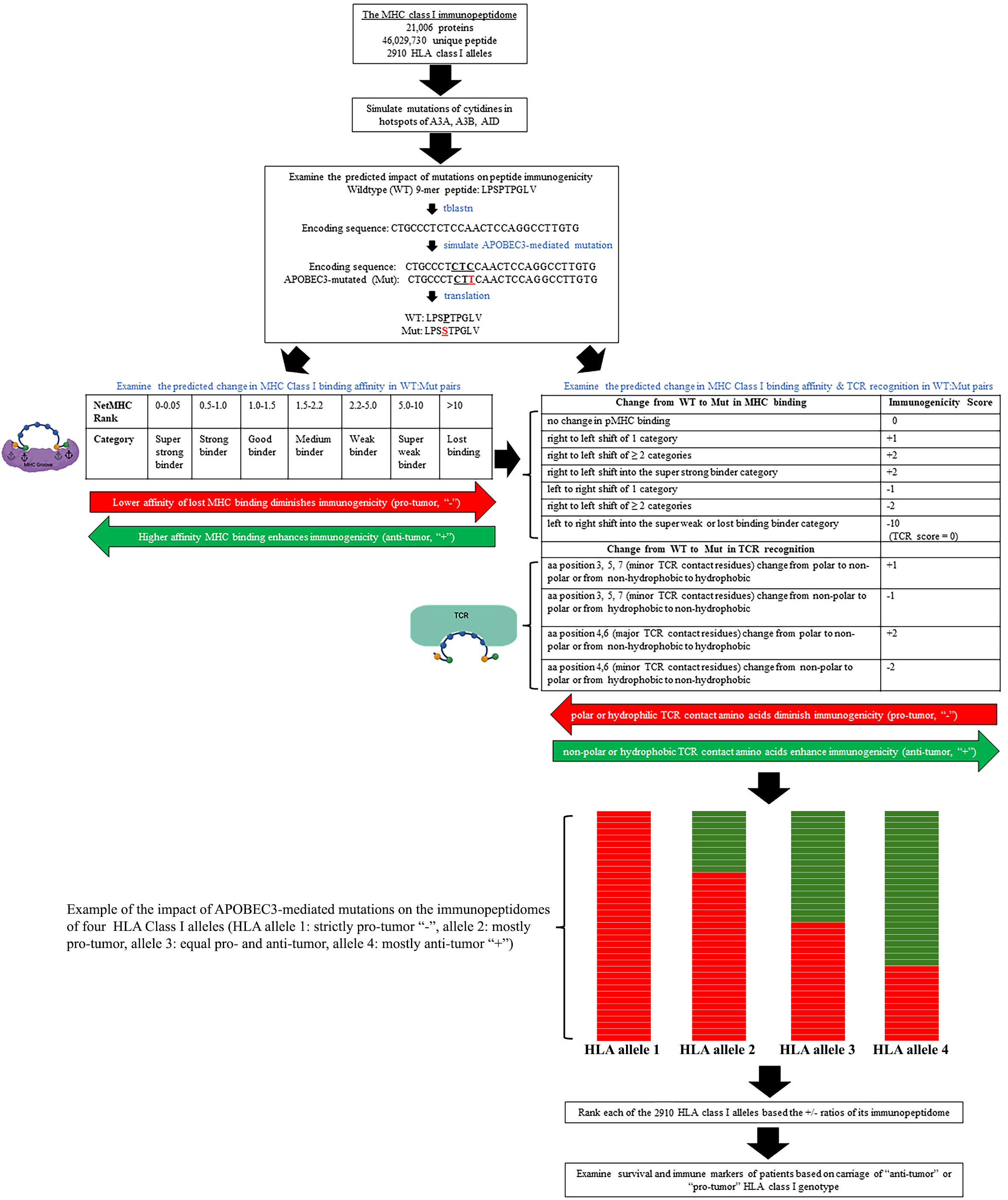
Prediction of the impact of APOBEC3-mediated mutations on the immunogenicity of the human immunopeptidome. The human MHC (HLA) class I immunopeptidome is comprised of millions of 8-11mer peptides that can potentially be bound and presented by several thousand human HLA class I molecules, making several billion peptide:MHC (pMHC) combinations that can potentially be presented for recognition by T cells. For each peptide epitope in the immunopeptidome, we recovered the genomic coding sequence and simulated mutations at favored hotspot motifs of APOBEC3A or APOBEC3B, and as control, AID. Following mutation simulation (scale described in table 1), we evaluated the predicted impact of each mutation event by comparing each wild-type (WT): mutant (MUT) pair, for the impact of the mutation on pMHC binding affinity (lower left), and pMHC + T cell receptor (TCR) binding affinity (lower right). For the pMHC analyses (lower left), we evaluated whether the percentile rank pMHC affinity was altered using NetMHC 4. We used two criteria to account for small inter-category movements: a percentile rank value change > 0.3 for a 1-category shift and a significant shift between the two extremes of a category (separated by distribution Z score > 0.5). Mutational events were thus binned into the following four categories “-“: lost or decreased immunogenicity=pro-tumour, “+”: gained immunogenicity=anti-tumour, NC: no change in immunogenicity, Synonymous: silent. The +/- ratio of +(enhanced immunogenicity=anti-tumor mutational events) / -(lost or diminished immunogenicity=pro-tumor mutational events) was thus calculated for the total mutations induced by each APOBEC3 enzyme, as well as for the immunopeptidome of each HLA allele. For TCR binding of the pMHC complex (lower right), we took into account the amino acid property changes in each WT:MUT pair which are predicted to impact TCR binding, in addition to the pMHC affinity changes. Changes in the hydrophobicity and polarity of amino acid substitutions at each position in each WT:MUT pair, from WT to MUT were determined using the Kyte-Doolittle and Grantham scales. To illustrate the magnitude of the changes, an index based on the quartiles of hydrophobicity and polarity substitution at each position was employed. Peptide positions 4 and 6 received a higher weight in score than positions 3,5,7, since the former are more dominant TCR contacts. In this manner, each mutational event was given a score that combines pMHC binding and TCR recognition. For the immunopeptidome of each HLA allele, the number of simulated mutation events in the pool of its restricted epitopes that resulted in a change in immunogenicity towards either a potential “+” (enhanced immunogenicity=anti-tumor mutational events) or “-“ (lost or diminished immunogenicity=pro-tumor mutational events) was tallied to obtain the probability of an APOBEC3-mediated mutation to instigate either an anti-tumor or pro-tumor event. The +/- ratio of + (anti-tumor mutational events)/- (pro-tumor mutational events) was thus calculated, considering changes in HLA and TCR binding affinities. In parallel to the mutation count of + and - events, the numerical scores which reflect the magnitude of changes in immunogenicity for pMHC and TCR binding for each mutational event were used to assess the magnitude of each change of immunogenicity event in either the “+” anti-tumor or “-“ pro-tumor direction. For each HLA allele, we then calculated a cumulative immunogenicity score (CIS) which is the sum of the individual mutational event’s score values for the HLA and TCR immunogenicity changes, in either the anti-tumor (+ values) or pro-tumor (- values) direction. The +/- ratio of these values indicates the anti-tumor/pro-tumor potential of each HLA allele, considering the number of mutational events that cause an immunogenicity change in either the anti- or pro-tumor directions, and also the magnitude of each immunogenicity change event as determined by the type and position of amino acid changes in the epitope. Four example HLA alleles are shown whose immunopeptidomes contain different proportions of + and – events. HLA alleles were then ranked based on the +/- ratio of their immunopeptidome after APOBEC3-mediated mutations.

**Table 1.** APOBEC3-mediated mutation simulation and prediction of immunogenicity consequences.

|  |  |
| --- | --- |
| **Starting wildtype human immunopeptidome** | The HLA class I immunopeptidome includes all unique 8-11mer peptides (~ 46 million) of which 28% (13,684,528) have a functional affinity (< 500 nM) for at least one of 2910 HLA class I alleles = ~ 1 billion peptide:MHC (pMHC) combinations |
| **Mutations simulated (C >T, G>A)** | APOBEC3A (CTC, <u>G</u>AG, TTC, <u>G</u>AA) |
|  | APOBEC3B (ATC, <u>G</u>AT, GTC, <u>G</u>AC) |
|  | AID (AG<u>C</u>, TAC, TGC, <u>G</u>CT, <u>G</u>TA, <u>G</u>CA) |
| **APOBEC3-mutable immunopeptidome** | 12,749,500 unique peptides=914,059,070 pMHC (93% of HLA class I immunopeptidome) |
| **Simulated mutation events** | APOBEC3A (1,430,782,669) |
|  | APOBEC3B (1057937974) |
|  | AID (2,013,086,955) |
|  | Total= 4,245,047,967 |
| **Genome coverage** | 381,534,562 nts (~ 12% of the genome) |
| **Total mutated pMHC combinations** | 6,009,308,016 |
| **Amino acid consequence of simulated mutations (pMHC)** | APOBEC3A non-synonymous 869,066,666 (61%) |
|  | synonymous 561,716,003 |
|  | APOBEC3B non-synonymous 571,472,553 (54%) |
|  | synonymous 486,465,421 |
|  | AID non-synonymous 1,320,733,062 (66 %) |
|  | synonymous 692,353,893 |

As shown in table 1, most peptide-encoding genomic sequences (93%) contained at least one AID/APOBEC3 hotspot. In total, ∼4.5 billion mutation events were simulated of which the majority (∼2.6 billion, 62%) yielded non-synonymous (NS) amino acid changes. A3B-mediated mutations had a lower likelihood of generating NS changes relative to A3A and AID (NS/S ratio of 1.2, 1.5 and 1.9, respectively), and mutations in A3B hotspots yielded less mutated peptides relative to A3A (∼571 vs 869 million) in total. The remaining epitopes devoid of APOBEC3 hotspots or in which mutations yielded synonymous codons may represent advantageous candidates for immunotherapy, being exempt from immune escape-inducing mutation by APOBEC3s.

As shown in Figure 1 (left panel), each APOBEC3-simulated mutation was analyzed for the change in the peptide:MHC (pMHC) binding affinity between that of the wildtype (WT) and the APOBEC3-mutated (Mut) pair, to the restricting HLA. Each WT-Mut pair was classified as having one of four outcomes: lowering/loss of pMHC binding affinity considered as a pro-tumour or immune escape potential (designated “-“), gain of pMHC affinity considered as anti-tumour or enhanced CTL recognition potential (designated “+”), no change in binding affinity “NC”, or synonymous mutations, the latter two being neutral events, neither pro- nor anti-tumor (Supplementary tables 1-3).

Across all WT-Mut pairs, A3A and AID were found to be 4.6 and 3.7 times more likely to produce “-” mutations than “+” mutations, indicating a stronger potential for promoting tumor immune escape. In contrast, A3B showed a less pronounced bias, with a 2.1-fold likelihood of generating “-” over “+”mutations. The median -/+ ratio across all HLA alleles was 20 for A3A, 19 for AID, and 7.7 for A3B, reflecting the lesser pro-tumor potential of A3B (Figure 2A).

**Figure 2.**
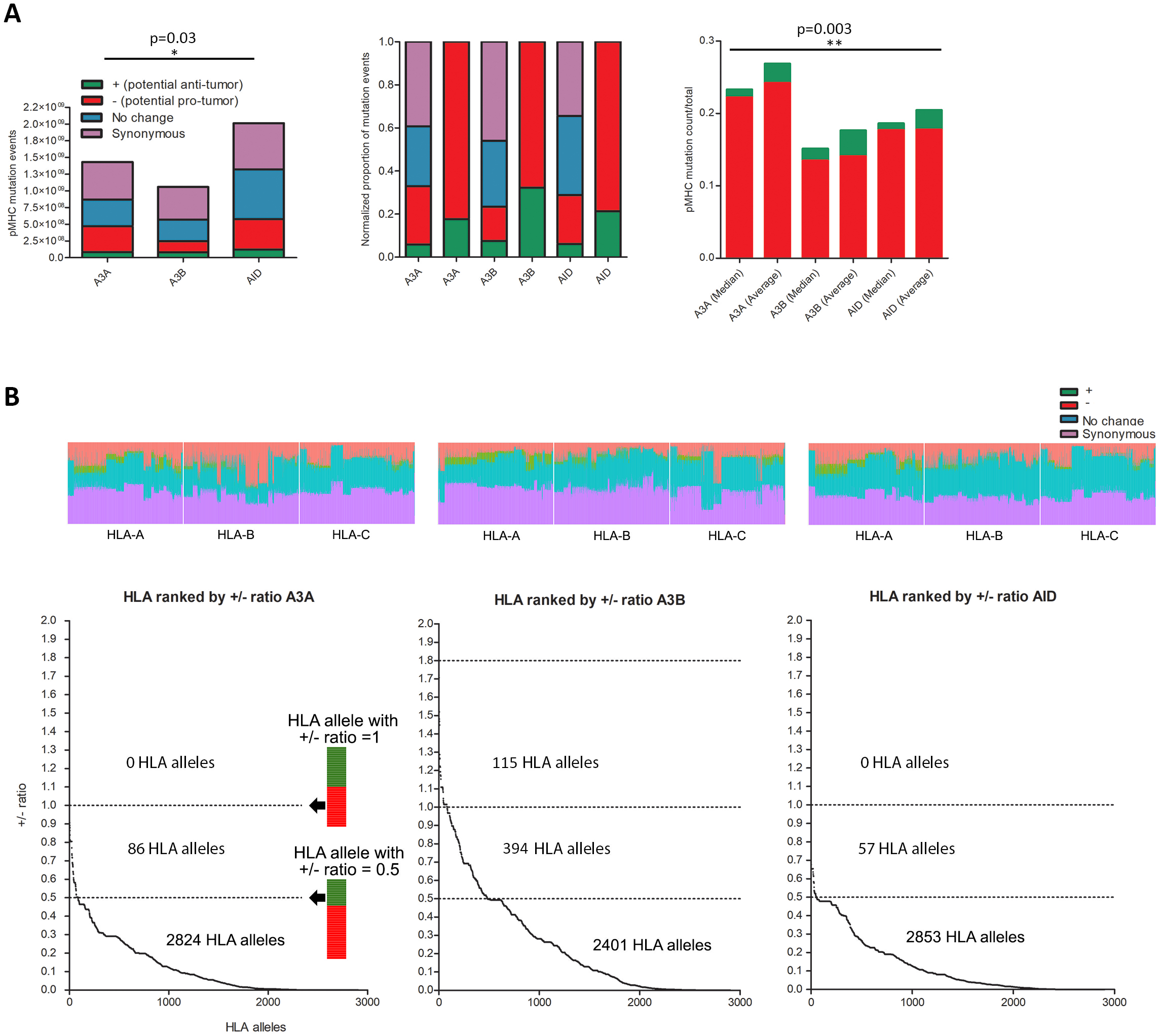
Impact of simulated APOBEC3-mediated mutations on changing the MHC binding affinity of the immunopeptidome. **A.** The left panel shows the total mutation events of each enzyme as broken down into the four categories of “-“: decreased pMHC binding = pro-tumour, “+”: enhanced pMHC binding = anti-tumour, “NC”: no change in pMHC binding affinity, and “synonymous”: silent mutations. The y-axis shows the raw count of mutational events. The middle panel shows the four category relative proportions normalized (1 = 100%) for each enzyme. The right panel shows the +/total and -/total mutational counts for each enzyme, as an average or median of the value determined for the immunopeptidomes of each of the 2910 HLA alleles. The “+” or “-“ mutational event counts was normalized to the total mutational count, for each of the 2910 HLA alleles, and the average and medians of this relative proportion across all HLA alleles is plotted. **B.** The top panel shows the proportion of the 4 categories for all 2910 HLA alleles, to demonstrate the variation among alleles. The bottom panel shows the 2910 HLA alleles ranked by +/- ratio of their immunopeptidomes as mutated by A3A, A3B and AID. For each HLA allele, the relative count of “+” (anti- tumour)/total MUT events was divided by “-“ (pro-tumour)/total MUT events to obtain a +/- ratio which is plotted for each of the 2910 HLA alleles.

Thus, while both A3A and A3B mutations predominantly induce non-synonymous changes in the immunopeptidome, A3A and AID mutations are more likely to lead to overall reduced peptide presentation potential by MHC class I

### Marked variation in HLA class I alleles influencing potential immune escape and neo epitope generation following APOBEC3-mediated mutation

We ranked HLA alleles by +/- ratio, reflecting how likely APOBEC3-mediated mutations of their restricted immunopeptidome was to generate enhanced “+” vs. diminished “-“ immunogenicity outcomes (Figure 2B, Supplementary tables 4-6). Among HLA alleles, a remarkable variation emerged in the +/- ratios (Figure 2B). The majority of HLA alleles fell at the extreme pro-tumor end as evident by two observations: first, A3A-, A3B- and AID-mediated mutations led to 100% pro-tumour “-“ outcomes (i.e. +/- ratio=0) for ∼15% of HLA alleles (428, 452, 411 alleles, respectively); second, A3A-, A3B-and AID-mediated mutations are 10-times more likely to yield pro- than anti-tumour pMHC outcomes (+/- ratio < 0.1) for 1820, 1330 and 1802 HLA alleles out of the total 2910 alleles, respectively. ∼500 HLA alleles were 1000-times more likely to support a pro-tumour outcome for either A3A-, A3B-, or AID-mediated mutations (+/- ratio < 0.001).

At the other end of the spectrum, a +/- ratio greater than 1 indicates an HLA allele whose immunopeptidome is more likely to have enhanced rather than diminished immunogenicity after APOBEC3-mediated mutation. A3A and AID mutations did not produce an a +/- ratio > 1 for any HLA allele, with less than 3% of alleles (86 and 57, respectively) showing a +/- ratio over 0.5, and the highest ratios being 0.9 and 0.7. In contrast, A3B mutations resulted in a significantly higher proportion of alleles (509, or 17%) with a ratio over 0.5, including 115 alleles with a ratio over 1, and the highest reaching approximately 2 (Figure 2B).

The data revealed two main findings. First, there is significant variation among HLA alleles in the degree to which their restricted immunopeptidomes’ immunogenicity is diminished or enhanced by APOBEC3-mediated mutations. Second, as shown in Figures 2B, comparing between A3A, A3B and AID, A3B-mediated mutations support the most anti-tumor outcomes (enhanced immunogenicity) relative to A3A and AID, both in the number of HLA alleles with higher +/- ratios and the higher value of the +/- ratios (Supplementary tables 4-6). Thus, most HLA alleles are skewed towards reduced immunogenicity of their restricted immunopeptidome, particularly under A3A and AID mutations; however, a subset of HLA alleles, especially under A3B-mediated mutagenesis, exhibits a higher likelihood of their restricted immunopeptidome gaining immunogenicity.

### Combined MHC binding and T cell receptor recognition score reveals that APOBEC3B- mediated mutations have the highest potential for neo-epitope formation

The immunogenicity of the CTL immunopeptidome not only requires pMHC formation, but also the recognition of the pMHC complex by the CTL through the T cell receptor (TCR). Thus, we sought to generate a more comprehensive immunogenicity score (CIS) for each WT:Mut peptide epitope pair, which includes T cell recognition in addition to pMHC formation which was evaluated above. To this end, we took into account the position and nature of amino acid changes in the TCR contact points (Figure 1, right panel).

For each HLA allele, +/- ratios were calculated in two ways: first, by CIS event, which consists of enumerating the number of mutational events that resulted in a negative (loss of immunogenicity) vs. positive (gain of immunogenicity) CIS change (mutational events with a - score or + score); and second, by CIS value, which includes the number of CIS events but also the qualitative weight of the CIS change in each negative or positive mutational event (the sum of all - scores or + scores, for the immunopeptidome of any given HLA allele) (Supplementary tables 7-9).

CIS event count analyses revealed that A3A- and AID-mutated immunopeptidomes predominantly resulted in reduced immunogenicity, with CIS event count -/+ ratios of 1.9 and 3.3-fold, respectively. In contrast, A3B-mediated mutations produced a more enhanced immunogenicity changes, with CIS event count -/+ ratios of approximately 1 (Figure 3A, left panel). The median +/- CIS count for A3A and AID were 0.6 and 0.2, while for A3B, it was 1.1, indicating more HLA alleles whose immunopeptidomes had higher +/- CIS ratios after A3B mutations (Figure 3A). As shown in Figure 3B, variation analyses among HLA alleles revealed that 453 HLA alleles had +/- CIS event and sum ratios >1 for A3A, while 1140 alleles had these ratios >1 for A3B. The AID- mutated immunopeptidome was the most pro-tumor, with only 70 alleles having a +/- CIS ratio >0.5, and none >1 (Figure 3B, C).

**Figure 3.**
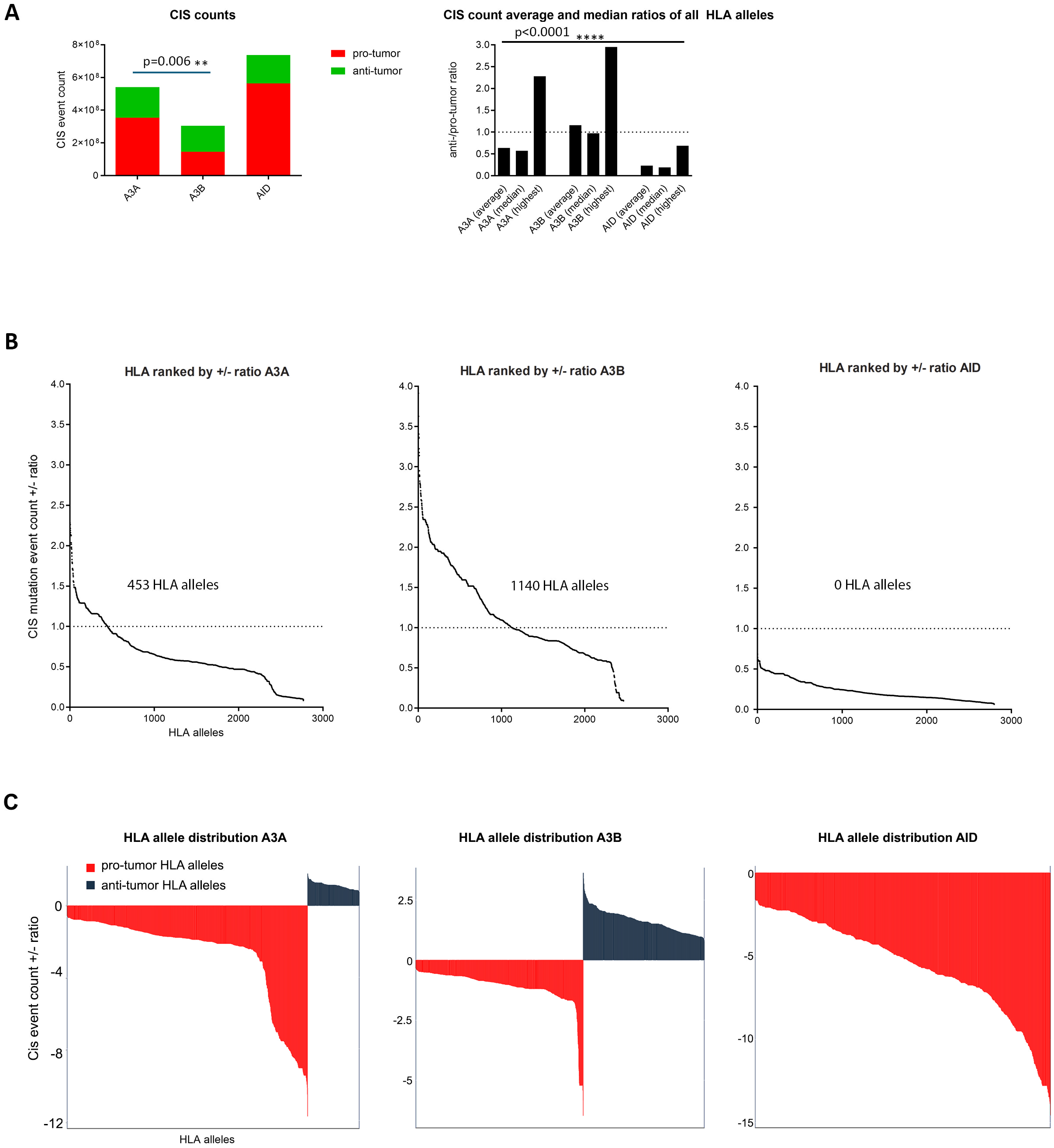
APOBEC3-mediated mutations alter peptide epitope immunogenicity by impacting pMHC binding and T cell receptor recognition. **A.** After ascribing a combined immunogenicity score (CIS) which accounts for changes in both pMHC binding affinity as well as TCR contact residues, for each mutational event, we tallied the count of APOBEC3-mediated mutational events that produced a “-“(diminished immunogenicity=pro-tumour potential) vs. “+” (enhanced immunogenicity=anti-tumour potential) in the immunopeptidome of each HLA allele. The anti-tumour “+” event counts were compared to the pro-tumour “-“ event counts for the total immunopeptidome as mutated by each enzyme (top left). The +/- ratio averages, medians and highest values of CIS even counts across the 2910 HLA alleles were derived (top right). **B.** For each of the 2910 HLA alleles, we tallied the count of CIS events (top row) in the enhanced immunogenicity “+“ and reduced immunogenicity “-“ directions and calculated a +/- ratio, based on which the HLA alleles were ranked. **C.** Based on the data plotted in 3B, a representation of variation across HLA alleles in the CIS event +/- ratios. +/- > 1 were considered “anti-tumor”, and +/- < 1 as “pro-tumor” HLA alleles.

The CIS value analyses revealed a similar pattern in that A3A- and AID-mutated immunopeptidomes showed a more reduced immunogenicity profile with CIS value -/+ ratios of 2.2 and 3.7-fold while A3B-mediated mutations produced a more enhanced immunogenicity profile with CIS sum -/+ ratios of approximately 1 (Figure 4A, left panel). Accordingly, the median CIS value sum for A3A and AID were 0.6 and 0.2, while for A3B, it was 1.1 (Figure 4A, right panel). As shown in Figure 4B, variation analyses across HLA alleles revealed that 453 and 461 HLA alleles had +/- CIS value sum ratios >1 for A3A, while 1029 alleles had ratios >1 for A3B. The AID-mutated immunopeptidome was the most pro-tumor, with 98 alleles having a +/- CIS ratio >0.5, and none >1 (Figure 4B, C).

**Figure 4.**
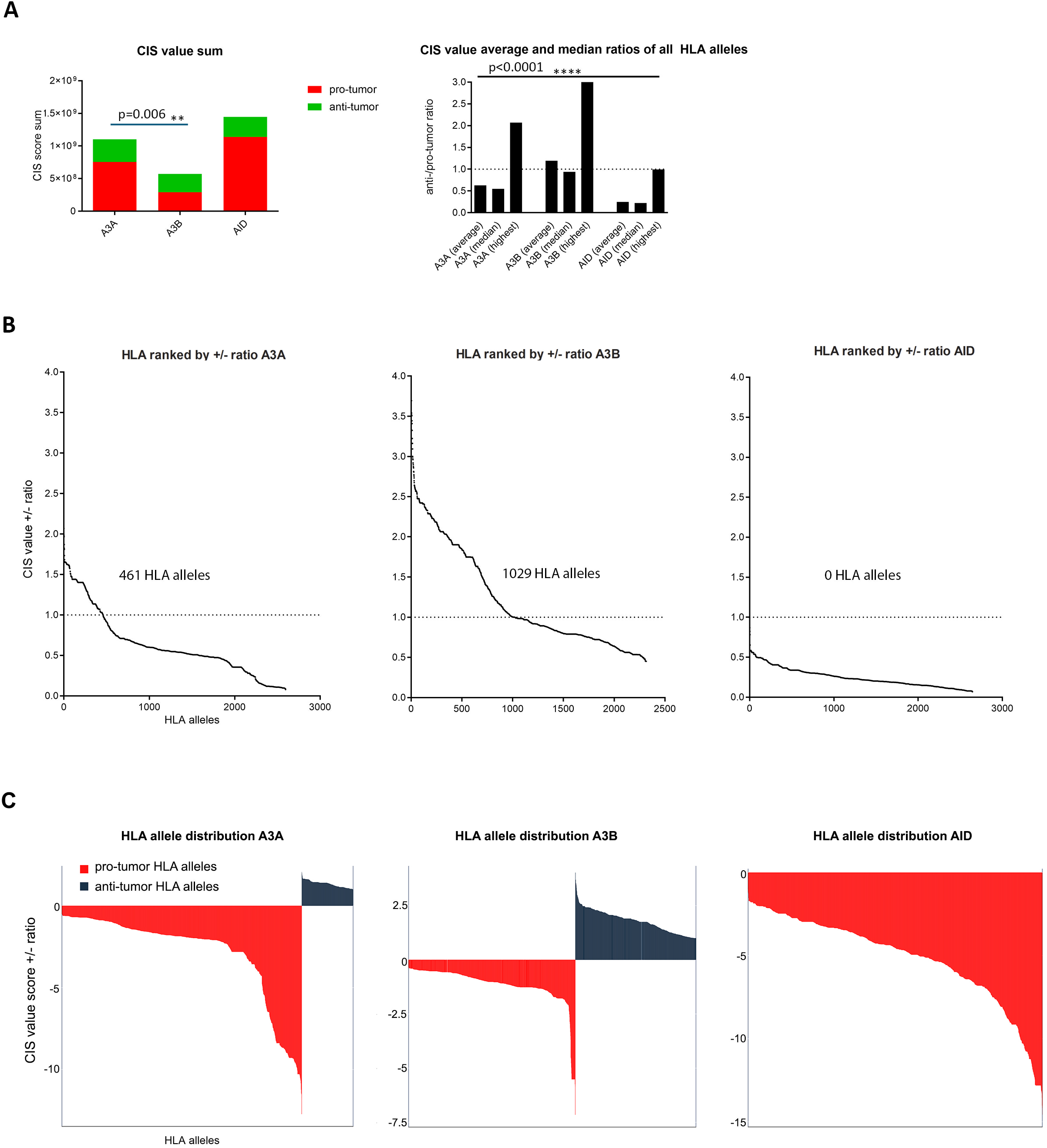
APOBEC3-mediated mutations have a differential impact on the magnitude of immunogenicity changes across HLA alleles. **A.** We calculated the sum of all “-“ value (diminished immunogenicity) scores, and. “+” value scores (enhanced immunogenicity) in the immunopeptidome of each HLA allele. The total anti-tumour CIS value of “+” for all HLA alleles was compared to the total pro-tumour CIS value “-“, representing an overall view of the whole immunopeptidome as mutated by each enzyme (top left). The +/- ratio averages, medians and highest CIS values across the 2910 HLA alleles were derived (top right). **B.** For each of the 2910 HLA alleles, we calculated a +/- ratio of CIS values and ranked HLA alleles. **C.** Based on the data plotted in 4B, a representation of variation across HLA alleles in the CIS value +/- ratios. +/- > 1 were considered “anti-tumor”, and +/- < 1 as “pro-tumor” HLA alleles.

As expected, the CIS event count and CIS value sum trends correlated linearly across HLA alleles, with variations due to APOBEC3 mutations showing a pattern of “+”>”-“>NC/synonymous, where the NC and synonymous categories showed the highest variation in the relative proportion of “+” changes (Supplementary Figure 2).

Hydrophobic and non-polar amino acids are prevalent in HLA-binding anchor terminal peptide positions and in the TCR contact positions of immunogenic CTL epitopes^18,19^. A3A-, A3B-, and AID-mediated mutations produced amino acid changes towards increased hydrophobicity and non-polarity in MHC anchor or TCR contacts, while A3A- and A3B-mediated mutations enhanced non-polarity compared to AID-mediated mutations at peptide anchor residues (Supplementary Figure 3). As a consequence of A3B-mediated mutations, 2 hydrophobic amino acids (L and M), 4 neutral amino acids (G, S, P, and H), and 2 hydrophilic amino acids (R and D) were modified to 6 hydrophobic (F, L, I, C, W, and Y), 3 neutral (K, N, and S), and 2 hydrophilic amino acids (E and G) (Supplementary Figure 4). A comparison of the pMHC and TCR components of CIS separately reveals that A3A- and A3B-mediated mutations generated a similar count and values of “+” TCR scores across the HLA alleles, but that A3B-mediated mutations led to significantly higher pMHC binding enhancements, approximately 3-fold better than A3A on average. (Supplementary Figure 5).

Thus, all three types of immunogenicity prediction analyses (pMHC, CIS event, CIS value) were in agreement in that A3B-mediated mutations have the highest potential to enhance CTL epitope immunogenicity compared to A3A and AID. This is reflected by a greater number of HLA alleles with positive CIS event and values for A3B, indicating a more favorable anti-tumor immune profile. APOBEC3-mediated mutations, especially from A3B, tend to increase hydrophobicity and non-polarity at MHC anchor and TCR contact positions in CTL epitopes. A3B-induced mutations notably enhance pMHC binding more than A3A-induced mutations.

### HLA class I genotype predicts survival in APOBEC3-mutated tumors

We generated a "triple index" to rank HLA alleles by combining each allele’s +/- rank across pMHC binding, CIS event count, and CIS value sum to identify HLA alleles whose immunopeptidomes have the highest +/- potential when mutated by A3A or A3B individually, or the combined action of A3A+A3B (Table 2, supplementary tables 7-11). We identified 332 HLA alleles for which the +/- ratio is >1 if mutated by A3A or A3B (Figure 5A, Table 2). Of these, 232 HLA alleles fall in the top 90^th^ percentile of the triple index rank for both A3A and A3B. These represent HLA alleles that have immunopeptidomes with a higher likelihood of + than – category mutations and therefore the highest potential to support enhanced anti-tumour CTL immunity, if a tumour’s genome is mutated by A3A, A3B or A3A+A3B. 55%, 54% and 60% of HLA alleles that supported the most anti-tumor “+” outcomes as a result of A3A-, A3B- and A3A+A3B-mediated mutations, respectively, are among the common, intermediate and well-documented (CIWD) list of HLA class I alleles (Table 2)^22^. We observed that the most anti-tumor HLA alleles (highest +/- ratios) for A3A-mediated mutations fell into the A1-A3 and A24 supertypes, for A3B-mediated mutations, into a broader range of supertypes including A1-A3, A24, B15, B18, B34, B44, B45 with the majority being in B44, and for A3A+A3B, a more restricted set of A1-A3 supertypes.

**Figure 5.**
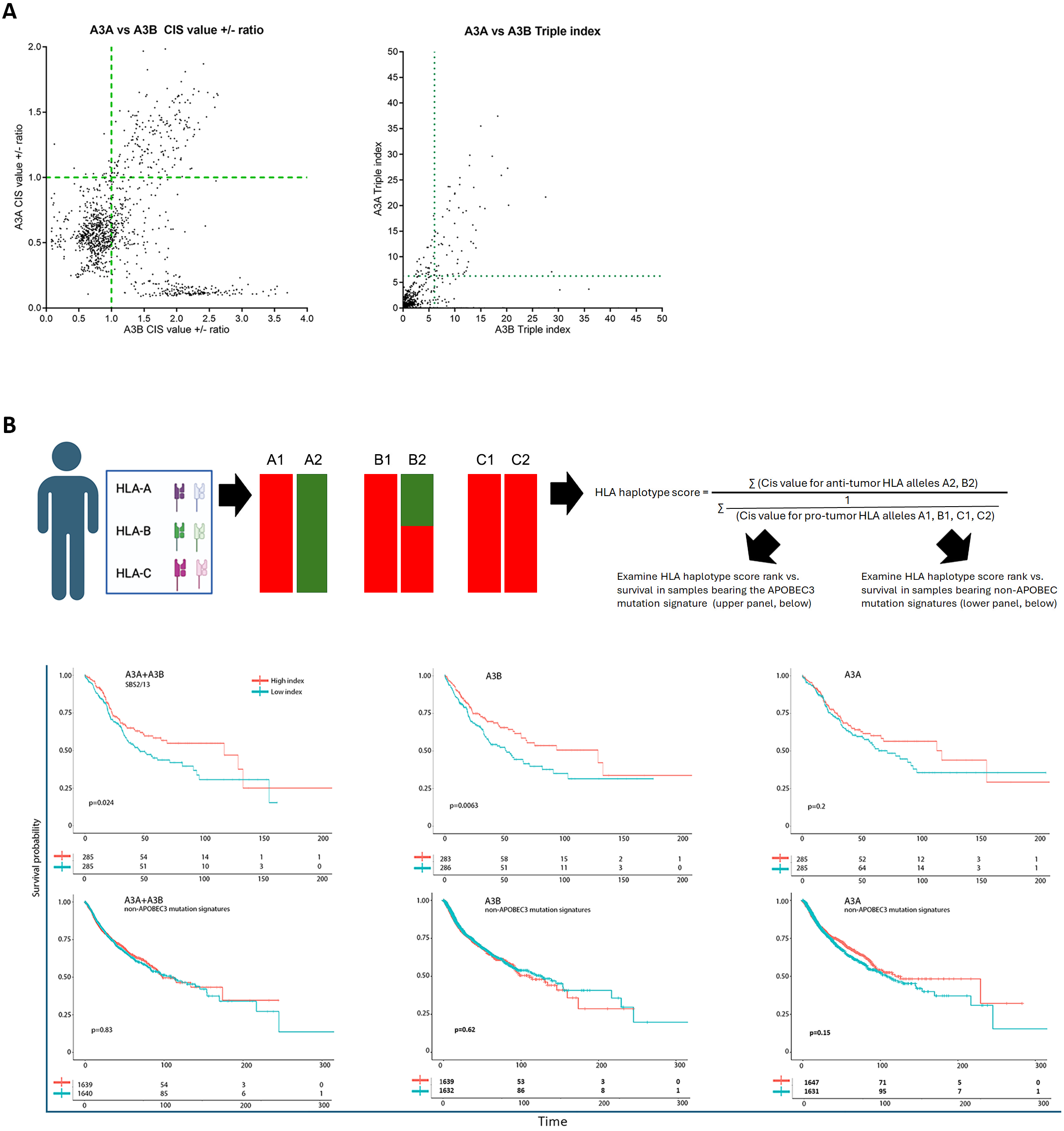
The HLA class I is a predictor of survival in APOBEC3-mutated tumors. **A.** The left panels shows the distribution of the 2910 HLA alleles based on CIS value +/- ratios of their A3A-mutated vs. A3B-mutated immuopeptidomes. The right panel shows the distribution of HLA alleles based on the triple index values of their A3A-mutated vs. A3B- mutated immunopeptidomes. The triple index integrates the +/- rank of each HLA allele relative to the average of all alleles, across the three measures of pMHC binding affinity, CIS event count, and CIS value. **B.** The top panel shows the scheme for deriving a personal HLA class I haplotype score for each person’s six HLA class I A, B and C alleles, from the individual HLA allele CIS values. Using the CIS value +/- ratios of each HLA allele, a combined +/- score for each individual in the TCGA dataset was devised which takes into account the +/- scores of each of the six HLA class I A, B and C alleles in an individual’s HLA haplotype. Patients were then ranked based on the cumulative +/- index of their HLA haplotype. The “High index” top third of the ranking represents patients that carry an HLA class I haplotype at the highest end of the cumulative +/- scores and therefore most potential for enhanced immunogenicity of the HLA class I immunopeptidome after APOBEC3- mediated mutations. At the other end of the spectrum, patients in the “Low index” fraction carry an HLA class I haplotype that falls at the lowest range of +/- scores, and therefore the most potential for lost or reduced immunogenicity and immune escape after APOBEC3- mediated mutations. Survival was then examined based for the “High index” vs. “Low index” patients for tumors that carry an APOBEC3 mutation signature (top row plots), and, as control, for those that do not carry an APOBEC3 mutational signature (bottom row plots). The left, middle and right panels denote different “High index” and “Low index” HLA haplotypes, based on the combined +/- scores of A3A+A3B, or A3B or A3A, alone.

**Table 2.**
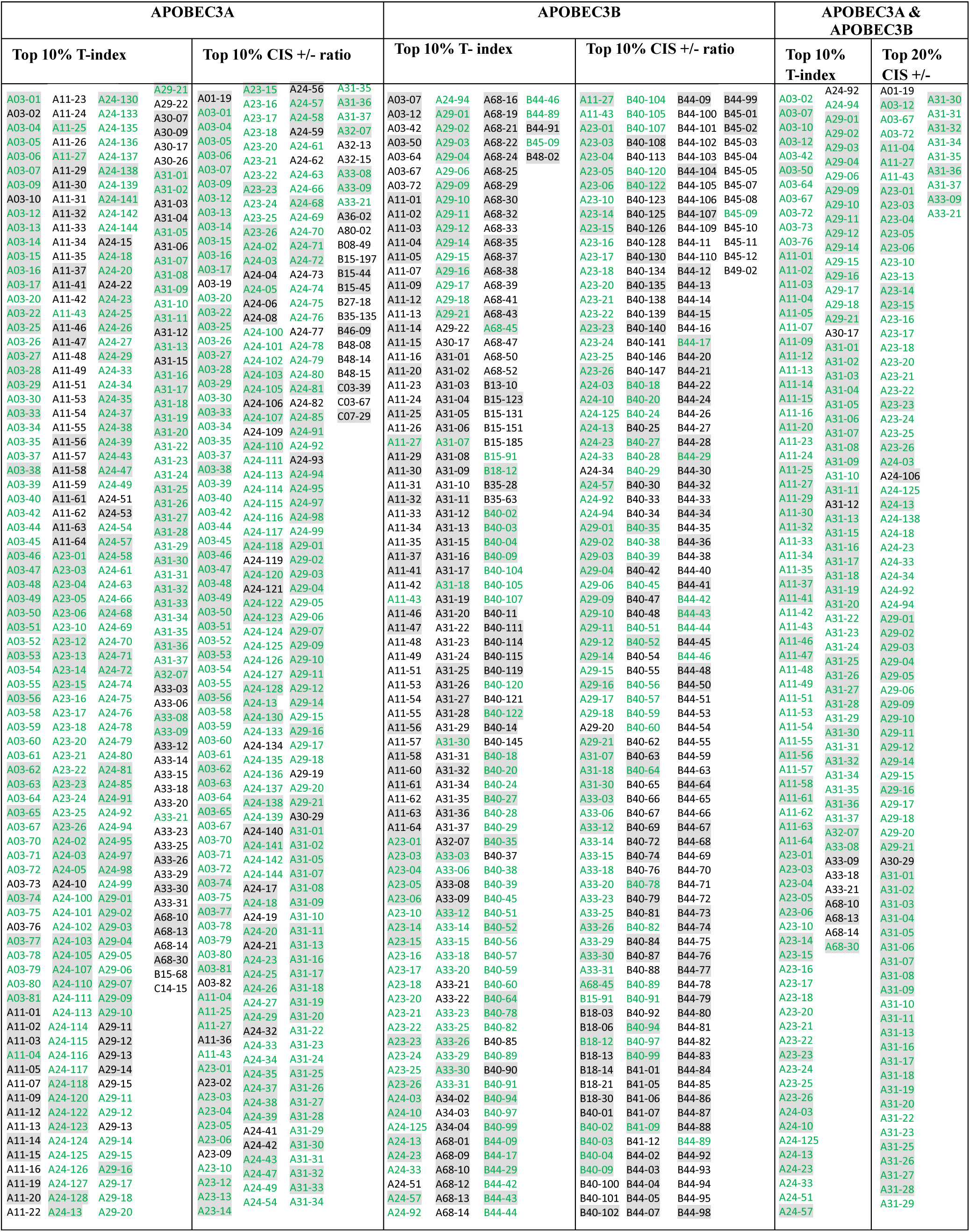
List of HLA alleles with the highest anti-tumor scores as a result of APOBEC3-mediated mutations of their immunopeptidome, according to the combined immunogenicity score (CIS), or triple index (TI) rankings. The HLA alleles colored green are shared across the top CIS and TI rankings, and alleles highlighted in gray are classified as common, intermediate or well documented (CIWD). The top percentile anti-tumor HLA alleles (highest +/- ratios) shown in this table for A3A-mediated mutations fell into the A1-A3 and A24 supertypes. For A3B-mediated mutations, a higher number and broader range of supertypes could support higher +/- ratios including A1-A3, A24, B15, B18, B34, B44, B45 with the majority being in B44. For A3A+A3B, a more restricted set of HLAs can support higher +/- ratios and these fell into the A1-A3 supertypes.

Taken together, the pMHC and pMHC-TCR (CIS) binding predictions demonstrate two concepts: first, a remarkable variation in the potential of APOBEC3-mediated mutations to diminish or enhance the immunogenicity of the CTL immunopeptidome dependent on HLA class I genotype; second, that, relative to A3A, mutations in A3B hotspots have a higher potential to enhance epitope immunogenicity, in the number of HLA alleles and the magnitude of the +/- ratio.

Our simulation data suggests that HLA class I alleles differentially channel APOBEC3-mediated mutations towards enhanced immunogenicity “+” or diminished immunogenicity “-“. If relevant, this would imply that carriage of specific HLA alleles would impact the anti-tumor CTL response, and specifically that carriage of HLA alleles with the highest +/- ratio might correlate with enhanced anti-tumor CTL immune response.

While our predictive analyses thus far revealed a +/- ratio for the immunopeptide of 2910 individual HLA alleles, each person carries six class I HLA A, B and C alleles in their haplotype. As shown in Figure 5B (top panel), we calculated a cumulative HLA haplotype +/- ratio for each patient in the TCGA, based on the combined value of their anti-tumor HLA alleles (defined as HLA alleles with CIS value +/- ratio > 1) divided by the combined value of their pro-tumor HLA alleles (defined as HLA alleles with CIS value +/- ratio <1). This calculation of an HLA haplotype score was carried out under three mutagenesis scenarios: A3A alone, A3B alone, or A3A+A3B, based on individual A3A- or A3B- induced mutagenesis CIS values, or a combined A3A+A3B CIS score (supplementary tables 7-11). Patients were ranked using this cumulative HLA genotype +/- index value, with “high” and “low” indexes denoting the anti-tumor and pro-tumor one-third of the spectrum. Thus, A3A+A3B “high index” patients have an HLA class I haplotype with the highest potential to channel A3A+A3B-mediated mutations towards enhanced immunogenicity, whereas A3A- alone or A3B-alone “high index” patients have an HLA class I genotype with the highest potential to channel A3A-mediated, or A3B-mediated mutations towards enhanced immunogenicity. We then sought correlations between this HLA haplotype index rank, and survival in the TCGA subset with the APOBEC3 mutational signature SBS2/13 which expectedly exhibit a higher expression level of A3A and A3B relative to tumors that do not bear the APOBEC3 mutational signature (Supplementary Figure 6).

Indeed, we observed a significant difference in survival in favor of the high index patients for A3A+A3B (p=0.024), and A3B (p=0.0063), and a non-significant trend in the same direction for A3A (p=0.2) (Figure 5B). As control, the same “high index” vs. “low index” HLA haplotype-based stratification did not yield survival analysis for patients whose tumors had non-APOBEC mutational signature (Figure 5B, bottom panel), indicating the specific role of HLA class I genotype in APOBEC3-mutated tumors.

Thus, the carriage of an HLA haplotype that includes HLA alleles that better channel APOBEC3- mediated mutations, particularly mediated by A3B, toward enhanced immunogenicity of the HLA class I-restricted immunopeptidome confers a survival advantage, specifically in APOBEC3- mutated tumors, and not in tumors bearing non-APOBEC3 mutational signatures. This highlights the combinatorial role of the mutational landscape and HLA class I genotype in shaping the survival outcome.

### Enhanced survival in APOBEC3-mutated tumors with high-index HLA class I correlates with markers of increased tumor-infiltrating activated CD8+ T Cells

Patients bearing high-index HLA class I haplotypes exhibit improved survival compared to low-index HLA class genotypes, due to the carriage of HLA class I alleles whose immunopeptidomes are more likely to become more immunogenic for CD8+ T cell recognition after APOBEC3- mediated mutations. This suggests that APOBEC3-mediated mutations enhance the CD8+ T cell response in high-index HLA class I patients.

We made three key observations supporting this model: first, we found that tumors with high-index HLA genotypes with the APOBEC3 mutational signature SBS2/13, had a higher presence of tumor-infiltrating CD8+ T cells compared to low-index HLA tumors (Figure 6A). This increased CD8+ T cell infiltration was observed precisely in tumors with survival differences (i.e. A3A+A3B and A3B mutational signatures) but not in tumors without the APOBEC3 signature (Figure 6A). Second, we carried out differential gene expression analyses on patients stratified based on high and low index HLA genotypes because of A3A+A3B- or A3B-mediated mutations, and whose tumors bear the APOBEC3 mutational signature. Indeed, the most substantial differences between high and low index HLA genotypes consisted of genes involved in CD8+ T cell-mediated tumor cell lysis (e.g., Granzymes, FasL, Perforin) and IFN-γ, a marker of an immune-active tumor microenvironment (Figure 6B). Third, differential expression analysis of genes that delineate subsets of neo-epitope reactive tumor-infiltrating CD8+ T cells ^23^ indicated increases in tumor-infiltrating resident memory (C4), memory effector (C7), and neo-epitope reactive tumor-infiltrating CD8+ T cells, in high-index HLA tumors with the APOBEC3 signature compared to low-index patients (Figure 6C). These findings collectively suggest that the improved survival of high-index patients is due to HLA class I alleles whose immunopeptidomes, when mutated by APOBEC3s, lead to a more robust anti-tumor CD8+ T cell response.

**Figure 6.**
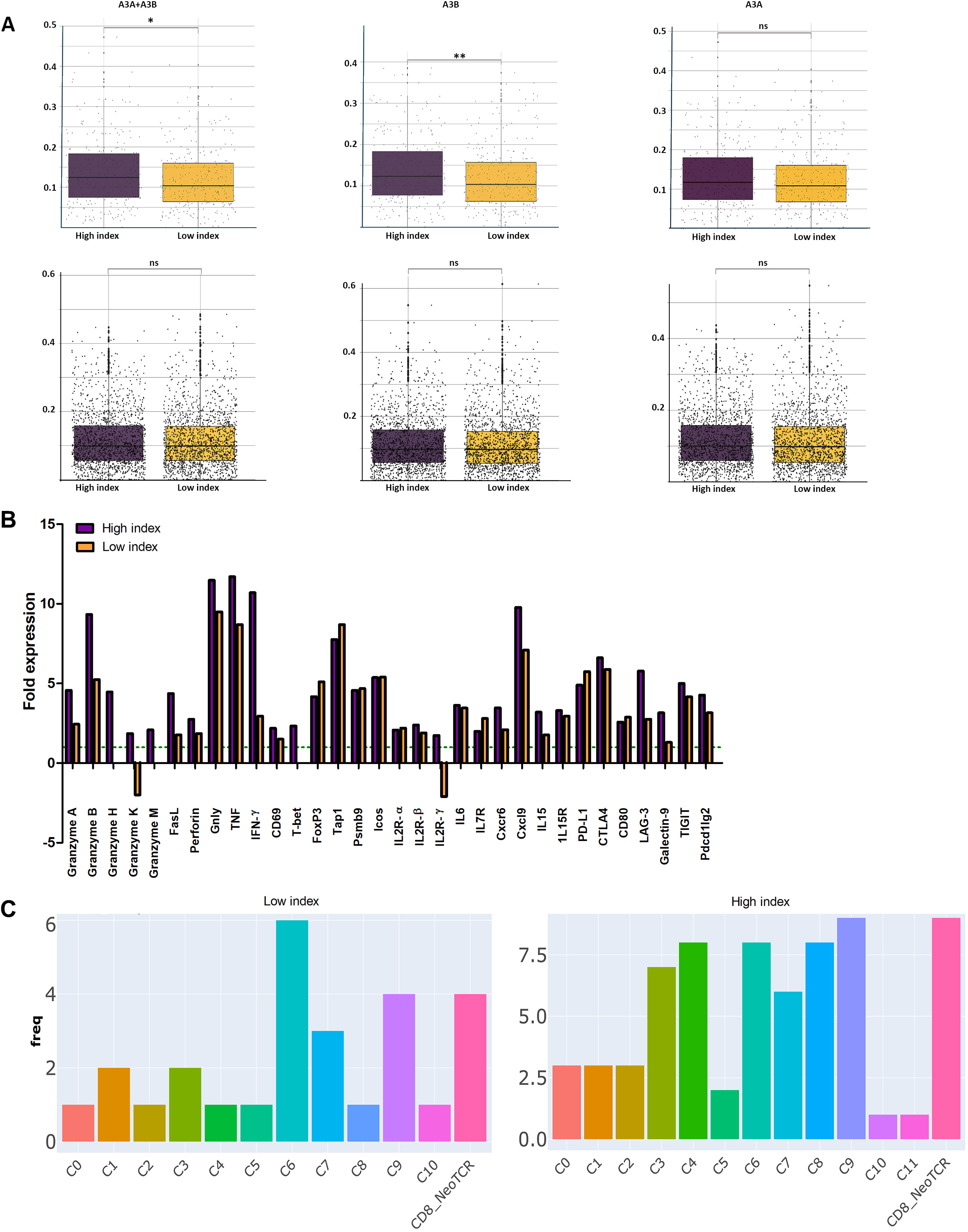
CD8+ T cell marker analyses of tumors stratified by HLA haplotype, APOBEC3 mutational signature, and survival. **A.** Tumors were stratified based on the carriage of High-index vs. Low-index HLA class I haplotype for A3A+A3B (left panel), A3B (middle panel), or A3A (right panel), and analyzed for CD8+ T cell infiltration according to immune landscape data^32^, and whether the tumors bore the APOBEC3 mutation signature (top panel), or non-APOBEC3 mutational signatures (bottom panel) **B.** Tumors were stratified based on the carriage of High-index or Low-index HLA class I haplotype for A3A+A3B, and the presence of APOBEC3 signature mutations, followed by differential gene expression analyses to examine expression levels of genes that are markers of CD8+ T cell activation (cytokines, transcription factors) and tumor killing by CD8+ T cells (granzyme, perforin, Fas Ligand) and T cell exhaustion (PD Ligand, Lag3, Icos). Differentially expressed CD8+ T cell markers between each set and tumors bearing non-APOBEC3 mutation signatures with Padj <0.05 were plotted. **C.** Differential gene expression data was analyzed for genes that delineate subsets of tumor-infiltrating T cells, including activated effector neoepitope-reactive CD8+ T cells^23^. APOBEC3-mutated tumors carrying high-index or low-index HLA class I haplotype for A3A+A3B were compared to tumors bearing non-APOBEC3 mutational signatures as baseline. Differentially expressed genes of > 2-fold and Padj <0.01 were considered as a filter, and the relative frequency of the filtered genes falling into each cluster was plotted.

## Discussion

Though the pro-tumorigenic roles of APOBEC3s’ in cancer initiation and progression are established from clinical and mouse model studies, a few studies have noted positive correlations between APOBEC3 expression and improved clinical outcomes^4,5,7,15–17,24,25^. In line with the known association between TMB and the success of immunotherapy, one anti-tumor activity of APOBEC3s could be neo-epitope generation. Such a context-dependent pro-tumor but also tumor-suppressive dual role including promotion of anti-tumor immunity has become increasingly established for A3B^15,25^. A3B-induced mutations lead to a library of neo-epitopes, which increase the efficacy of immune inhibitor blockade therapy intermediated by CTL and NK cells. A3B activity also induces anti-tumor immune responses and CD4^+^ T cell-mediated antigen-specific tumor growth inhibition^15^. A3B-mediated mutagenesis sensitizes HER2-driven mammary tumors to anti-CTLA-4 monotherapy and CTLA-4/anti-PD1 combination therapy^25^. Conversely, APOBEC3-mediated mutations could generate immune escape mutations. In clinical and mouse model studies, immune evasion mutations would be over-represented compared to those that boost immune recognition. The predictive phase of our work was motivated by the need to decipher the balance of APOBEC3’s potential to generate immune escape vs. immune boosting mutations in an unbiased sandbox.

We report several observations in support of a clinically significant role for APOBEC3-mediated mutations in regulating the anti-tumor CTL response: first, that relative to A3A, A3B-mediated mutations have the highest potential to enhance the immunogenicity of the CTL immunopeptidome, in agreement with the aforementioned studies on the anti-tumor roles of A3B in neoepitope generation. Second, there is an infinite level of variation among HLA class I alleles: some alleles have immunopeptidomes that when mutated by APOBEC3s support entirely a pro-tumor immune escape consequence, while other alleles have immunopeptidomes with a higher potential for generation of neo-antigens than escape variants. Third, the carriage of the predicted pro- or anti-tumor HLA class I genotypes, as a cumulative function of one’s HLA haplotype correlate with patient survival and CTL tumor infiltration.

Many of the HLA alleles whose immunopeptidomes generated the highest +/- values (anti-tumor potential of enhanced immunogenicity) when mutated by A3A and/or A3B, and the carriage of which correlates with survival, are among the CIWD HLA subset. Several belong to HLA-A03, HLA-A24 and HLA-A01A24 supertypes. HLA-A*03 is a negative biomarker for patients undergoing immune checkpoint inhibitors therapy^21^. Based on our data, we speculate that the patients who carry HLA-A*03 with high APOBEC3 levels would have an abundance of neo- antigens leading to preferential elimination of APOBEC3+ tumors. Given the correlation between expression of APOBEC3 and immune checkpoint molecules, these would also be less sensitive to checkpoint inhibition therapy.

This work highlights the importance of considering an individual’s HLA genotype in APOBEC3- mutated tumors. HLA genotype can serve as a biomarker to guide immunotherapy and future APOBEC3 inhibition therapeutic approaches^26^. Study limitations include not assigning differential weights in mutation simulation for secondary DNA structure which may generate regions favored by each APOBEC3 enzymes. Including this level of refinement would present a challenge as these regions would differ across individual tumors. Second, it is possible that the loss of immunogenicity of an epitope due to APOBEC3-mediated mutations could coincide with gain of binding to another co-inherited HLA. In addition, within a given protein, an entirely new epitope can emerge because of APOBEC3-mediated mutations. These questions can be addressed at the individual genomic level.

## Conclusions

This work is the first to conduct a genome-wide scan quantifying the impact of A3A- and A3B- mediated mutations on immunogenicity changes in the human immunopeptidome across thousands of HLA Class I alleles. APOBEC3-mediated mutations can lead to either neo-epitope generation or immune-escape epitopes, respectively resulting in enhanced or diminished CD8+ cytotoxic T cell (CTL) responses. We found that these two opposing consequences of APOBEC3- mediated mutations are not random, but rather vary markedly across the immunopeptidomes of different HLA Class I alleles. Accordingly, we found that HLA genotype serves as a predictor of survival and CTL activation, specifically in APOBEC3-mutated tumors. This novel prognostic role of HLA class I in APOBEC3-mutated tumors indicates that the impact of APOBEC3-mediated mutations in shaping the CTL immunopeptidome is significant enough to influence survival. Furthermore, identifying HLA genotype as a biomarker in APOBEC3-mutated tumors offers a promising avenue for personalized immunotherapy and potential A3 inhibition strategies.

## Methods

### Scoring the genome-wide impact of APOBEC3-mediated mutations on the CTL immunopeptidome

Figure 1 shows an overall scheme of the workflow and Table 1 contains information on the scale of immunogenicity predictions. The reference human HLA class I-binding immunopeptidome includes all possible 8-11mer peptides from ∼ 21,000 protein including 71,000 isoform sequences, and ∼46,000,000 unique peptides, whose HLA restriction includes 2,910 HLA class I alleles. This dataset (https://zenodo.org/record/1453418#.YoPo2ejMLUQ) which includes a total of 987,968,036 peptide:MHC (pMHC) combinations formed the initial basis of our AID/APOBEC3- mediated mutation simulation (Table 1) ^27^. tBlastn was used to find the corresponding encoding DNA coding sequence, using the following parameters: Database (Reference RNA sequences (refseq-rna)), Homo sapiens (taxid:9606), Max Target (100), Expected threshold (10,000), Word size (3), Low complexity filter (deselected). The encoding DNA sequence, including 2 nucleotides preceding the first codon were subject to mutation simulated at APOBEC3 hotspots (YTC (A3A), RTC (A3B), and WRC (AID), Y= purine, R= pyrimidine, W=A/T) ^28^ as follows: (AID: "TGC", "TGT"),(AID: "AGC", "AGT"),(AID: "TAC", "TAT"),(AID: "GCA", "ACA"),(AID:"GTA", "ata"),(AID:"gct", "act"),(A3A:"ttc", "ttt"),(A3A: "gaa", "aaa"),(A3A: "ctc", "ctt"),(A3A: "gag", "aag"),(A3B:"atc", "att"),(A3B:"gat", "aat"),(A3B: "gtc", "gtt"), (A3B:"gac", "aac").

NetMHCpan (version 4.1) was used to predict pMHC binding affinities changes between the wildtype (WT) and APOBEC3-mutated (Mut) peptides, which were categorized as shown in Figure 1. In addition, we predicted the immunogenicity impact of changes in the TCR binding portion of each peptide, as shown in Figure 1. This combined immunogenicity score (CIS) was obtained by considering amino acid hydrophobicity, polarity changes at the terminal MHC-binding anchor residues (1,2,9,10) and TCR contact positions (residues 4 and 6 received a higher weight than 3,5, and 7, since they are typically the more dominant TCR contacts) ^29,30^. For each HLA allele, we tallied the counts of WT to Mut changes (CIS event count), as well as the magnitude of WT to Mut changes (CIS value) that were predicted to decrease immunogenicity (“-“, pro-tumour) increase immunogenicity (“+”, anti-tumour) and calculated a +/- ratio. The 2910 HLA alleles were ranked by their +/- ratios of pMHC binding, CIS event count, and CIS values.

### HLA genotype- and APOBEC3-based solid tumor survival analysis

We used mutSignatures (version 2.1.5)^31^ for extraction and analysis of mutational signatures in 8956 sequenced tumors from TCGA encompassing 30 cancer types and HLA OptiType as well as clinical data^31,32^. The reliability of de novo extracted signatures was assessed by inspecting the silhouette plot at the end of the signature extraction process. The extracted signatures were then compared and matched with the COSMIC database to identify the cases bearing the COSMIC signatures 2 and 13 of APOBEC3s.

Figure 5B shows the scoring system developed to evaluate the likelihood of an individual’s HLA class I haplotype (six A, B, C alleles) to support the balance of pro-tumor or anti-tumor immunity because of APOBEC3-mediated mutations. To determine anti- or pro-tumor potential, if the anti-tumor (+)/pro-tumor (-) CIS ratio was > 1 or <1, an HLA allele was considered as anti- or pro-tumour, respectively. An HLA allele with a +/- CIS ratio of 1.6, has a restricted immunopeptidome that is 1.6 times more likely to result in an anti-tumor than a pro-tumor outcome when mutated by A3A or A3B. Conversely, an HLA allele with a +/- CIS ratio of 0.1, has a 10-fold higher likelihood of supporting a pro-tumour decreased immunogenicity outcome when its immunopeptidome is mutated by APOBEC3, and was thus assigned a pro-tumor value of 1/CIS=10. To determine an overall score for each patient, we first determined which of six HLA A, B and C alleles of an individual’s haplotype fell into the anti-tumor vs. pro-tumor category, as described by CIS +/- ratio. Then, a cumulative +/- ratio for each individual was then derived using the individual’s cumulative HLA anti-tumor CIS values as the numerator, and cumulative HLA pro-tumor 1/CIS values as the denominator, according to the following formula, with a specific example shown in Figure 5B:

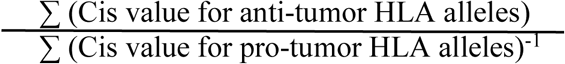

This cumulative +/- ratio reflects the combined potential of the immunopeptidomes of each individual’s six A, B and C Class I HLA alleles (HLA haplotype) towards either enhanced or diminished immunogenicity due to A3A-, A3B- or A3A+A3B-induced mutations. Thus, the higher an individual’s cumulative haplotype +/- ratio, the higher is their immunopeptidome’s potential for enhanced immunogenicity due to APOBEC3-mediated mutations. Based on this +/- index, patients were ranked, with "high index" or "low index" groups respectively denoting the top (HLA haplotypes with the highest or most anti-tumor-biased +/- values), or bottom (HLA haplotypes with the lowest or most pro-tumor-biased +/- values) one third. Using the CIS values derived from A3A-, A3B-, or A3A+A3B-mediated mutations (Supplementary tables 7-11), patients were ranked for a +/- ratio index for each of the three mutagenesis scenarios.

Correlations were then sought between the +/- ratio index and survival using linear models, and the survival (version 3.3.1) and survminer (version 0.4.9) packages in R^29^. The primary query of interest was overall survival, defined as the time from diagnosis to the last follow-up for which the Kaplan-Meier estimator was used. Survival curves were generated, and log-rank tests were conducted to compare survival curves between different groups.

### Immune marker and differential gene expression analyses

To identify differentially expressed genes and conduct immune marker analyses, we utilized DESeq2 and tumor immune landscape data as previously described^23,32^. TCGA sample count data were downloaded and used as the input. For A3A, A3B, as well as A3A+A3B combined we defined high and low index quantile groups. The count matrix was imported into R and converted into a DESeq2 dataset object. The APOEBC3 mutational signature and A3A and A3B TPM values were used as covariates in the final design model. P-values obtained from the differential expression analysis were adjusted for multiple testing using the Benjamini-Hochberg procedure to control the false discovery rate (FDR). Genes with an adjusted p-value (Padj) <0.05 and an absolute log2 fold change >1 were considered significantly differentially expressed.

### Statistical analysis

For the predictive phase of the work on differences among the APOBEC3 enzymes, linear models were used to determine statistical significance. Wilcoxon test was used to compare means. For survival analyses, Kaplan Meyer log-rank tests were conducted to compare survival curves between different groups.

## Data availability

Data are available at https://github.com/larijani-lab/

## Acknowledgments and Funding

The survival and immune biomarker results presented here are based upon data generated by The Cancer Genome Atlas (TCGA) via dbGaP (accession number phs000178.v11.p8), funded by the National Cancer Institute (NCI) at the National Institutes of Health (NIH).

Supplementary Raw Data Tables: HLA allele immunogenicity prediction scores and rankings*: https://github.com/larijani-lab/Supplementary-Files-

## List

Supplementary table 1: pMHC counts A3A

Supplementary table 2: pMHC counts A3B

Supplementary table 3: pMHC counts AID

Supplementary table 4: pMHC ranked A3A

Supplementary table 5: pMHC ranked A3B

Supplementary table 6: pMHC ranked AID

Supplementary table 7: CIS count, CIS value, Triple index A3A

Supplementary table 8: CIS count, CIS value, Triple index A3B

Supplementary table 9: CIS count, CIS value, Triple index AID

Supplementary table 10: Top anti-tumor HLA list combined A3A+A3B

Supplementary table 11: Top pro-tumor HLA list combined A3A+A3B

**\*** (data files too large to include in the manuscript file)

---

**Supplementary Figure 1.**
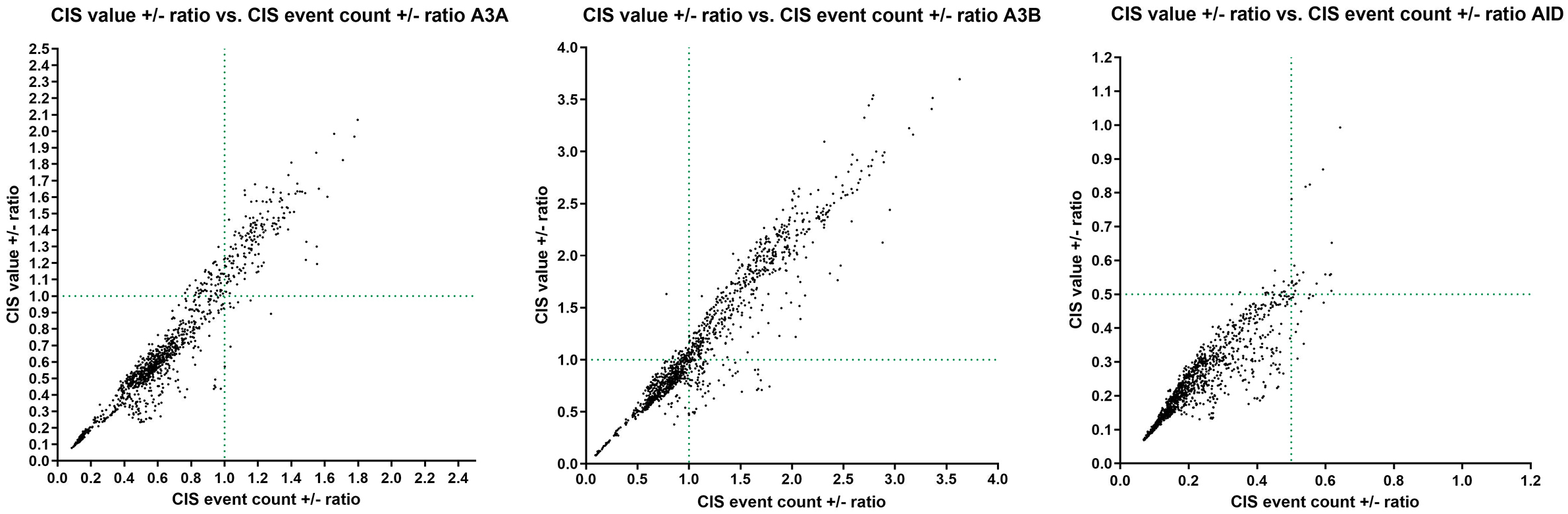
HLA allele distribution by CIS value +/- ratio relative to CIS event count +/- ratio. For A3A, AID and A3B, the CIS event count. Vs. CIS value +/- ratios were plotted with lines at a ratio of 1 in order to separate the HLA alleles with the highest +/- ratios across both metrics (upper right quadrant). The data are derived from appendix tables 7-9. For all three enzymes, a linear correlation was observed between HLA alleles distributed based on CIS event count vs. CIS value. This indicates that, as expected, the HLA alleles whose restricted peptidomes generated more events in the anti- or pro-tumor direction considering both pMHC and TCR recognition, also produced qualitatively larger shifts in either the pro- or anti-tumor direction. Since for AID, there were no HLAs with anti-/pro-tumor ratios > 1, 0.5 was chosen to delineate the HLA allele subset with the highest anti-tumor potential with respect to CIS value and CIS event count. For A3A and A3B, we observed 343 and 627 HLA alleles, respectively, with both the CIS event count and CIS value +/- ratio > 1, thus having a peptidome, that is more likely to generate an anti-tumor outcome when mutated by the respective enzyme. For AID, we observed 50 HLA alleles with both ratios > 0.5. For all three enzymes, we also observed outlier HLA alleles that had ratios that were >1 (for A3A and A3B) or > 0.5 (for AID) for either the CIS event count, or the CIS value, but not for both, representing the upper left and lower right quadrants. Among these, there were more HLA alleles in the lower right quadrants, representing alleles with a relatively high propensity for generation of more anti-tumor than pro-tumor epitopes but for which, the relative magnitude of the shift of the pMHC:TCR complex shift towards the anti-tumor outcome as a result of APOBEC3-mediated mutation was relatively more modest. In total, there were 125, 213, and 50 such alleles for A3A, A3B and AID.

**Supplementary Figure 2.**
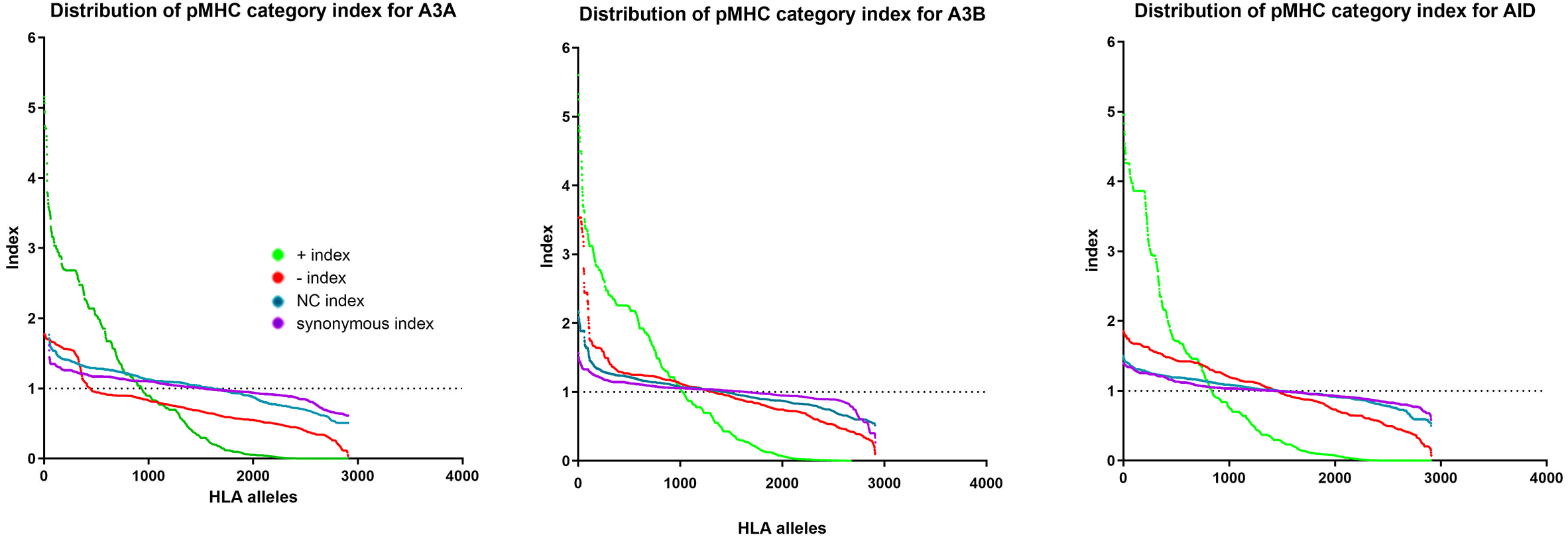
Distribution relationships among the four categories of pMHC binding evaluation due to APOBEC3-mediated mutations. Comparison of variability across HLA alleles in each of the four categories. For each category, the event count/total mut event count was derived. The ratios for each category were averaged across the 2910 HLA alleles, and then each HLA’s value was divided by the average to obtain an index value. Thus, an index value above 1, indicates that an HLA’s value was above average. The index range and distribution pattern were used to quantify the variation in each of the four event types. The level of variation among HLA alleles as a result of APOBEC3- mediated mutations followed the pattern of “+”>”-“>NC/synonymous. That is, the two categories that are inert with respect to pMHC binding affinity changes (NC, and synonymous) exhibited the least variation among HLA alleles, combined together, only varying between 20-70% of the total mutational events. The highest level of variation among HLA alleles was in the relative proportion of the “+” category changes (infinite range from 5-fold to zero), reflecting our observation that in general the extreme pro-tumor HLA alleles gained a +/- ratio of (near) zero due to having very few or no mutations in the “+” anti-tumor direction rather than having a significantly higher count of “-“ pro-tumor mutations.

**Supplementary Figure 3.**
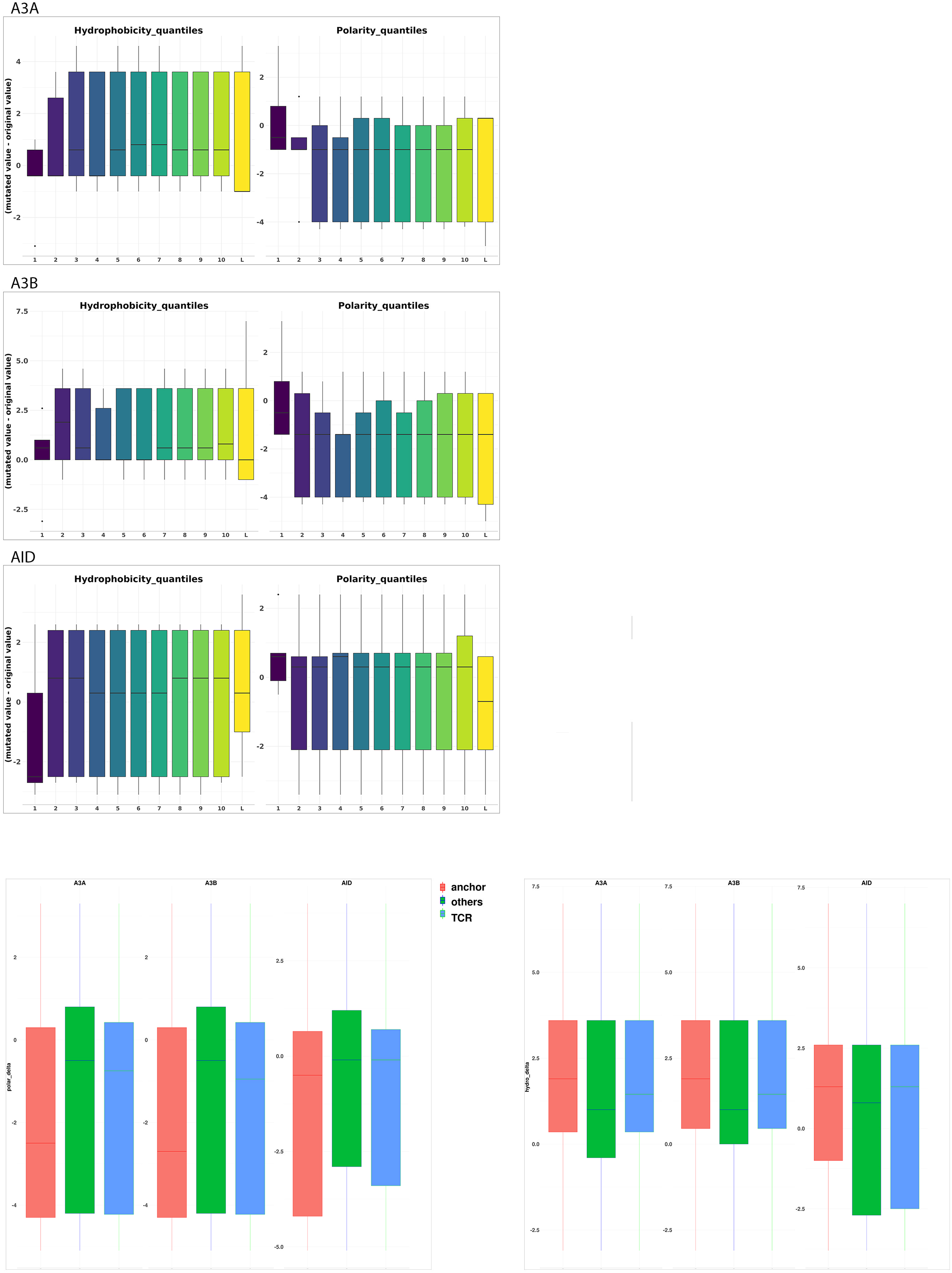
Changes in amino acid properties of peptide epitopes at HLA binding anchor residues and TCR contact positions due to APOBEC3-mediated mutations. To explain the changes in immunogenicity, non-synonymous APOBEC3-driven mutations were examined for specific changes in amino acids at positions along the length of the 8-11mer peptide epitopes including the end position (HLA anchor) and middle positions (TCR contacts). For each WT to MUT epitope mutation event, the hydrophobicity and polarity of amino acid substitutions at each position in the WT and MUT sequences were determined using the Kyte-Doolittle and Grantham scales, respectively. To illustrate the magnitude of the changes, an index based on the quartiles of hydrophobicity and polarity substitution at each position was employed, and delta values represent the change from WT to MUT. The top panel demonstrates the consequences mutations mediated by each of the APOBEC3 enzymes at each amino acid residue along the length of the 8-11mer peptides, and the bottom panel represents the cumulative changes an HLA anchor vs. TCR contact vs. the rest of the amino acid positions.

**Supplementary Figure 4.**
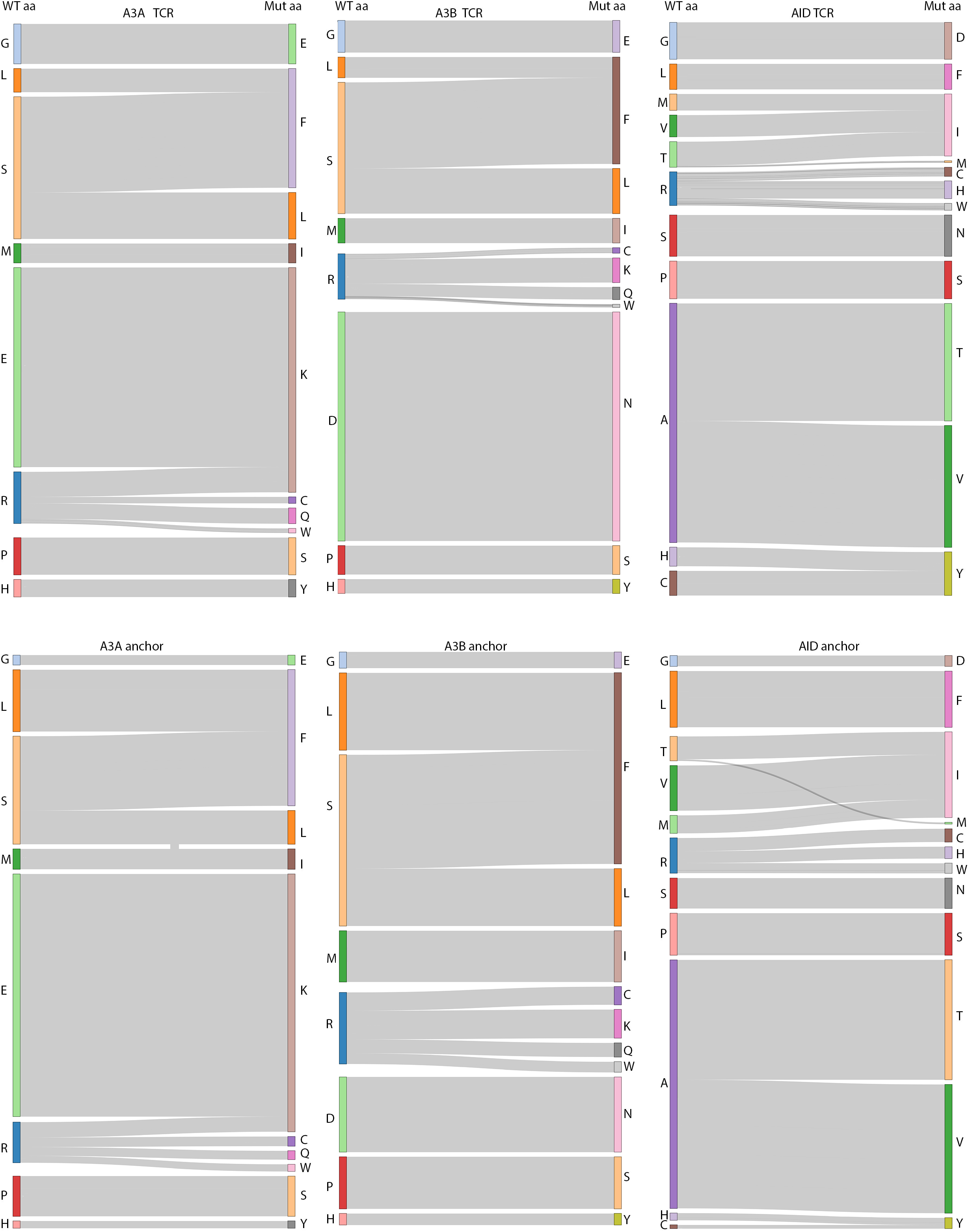
Specific amino acid change consequences of APOBEC3-mediated mutations in peptide epitopes. Sankey plots demonstrate the specific amino acid alterations and frequences as a result of mutations mediated by each APOBEC3 enzyme. The top panel shows changes at TCR contact middle positions in the 8-11mers, and the bottom panel shows changes in the HLA binding anchor residues at terminal positions.

**Supplementary Figure 5.**
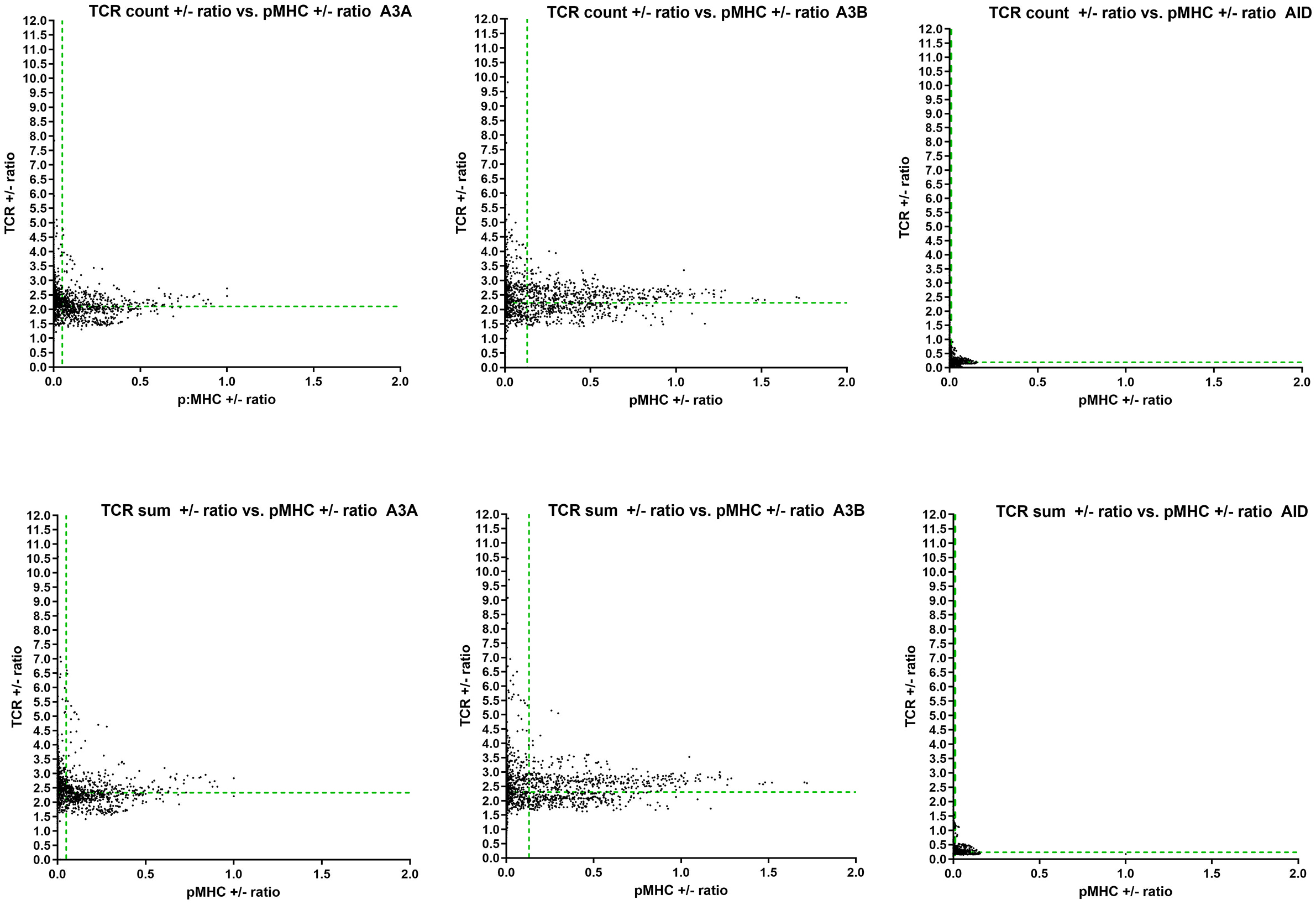
Comparison of the impact of the pMHC vs. the TCR recognition components of the CIS value changes. The cumulative immunogenicity index (CIS) takes into account how APOBEC3-driven mutations impact immunogenicity combining the two halves of the T cell epitope recognition equation: pMHC binding as well as pMHC-TCR recognition. To determine if mutations mediated by each APOBEC3 enzyme impact CIS differentially due to altering pMHC binding affinities vs. TCR contact positions, the two scores were separately considered and plotted for all 2910 HLA alleles.

**Supplementary Figure 6.**
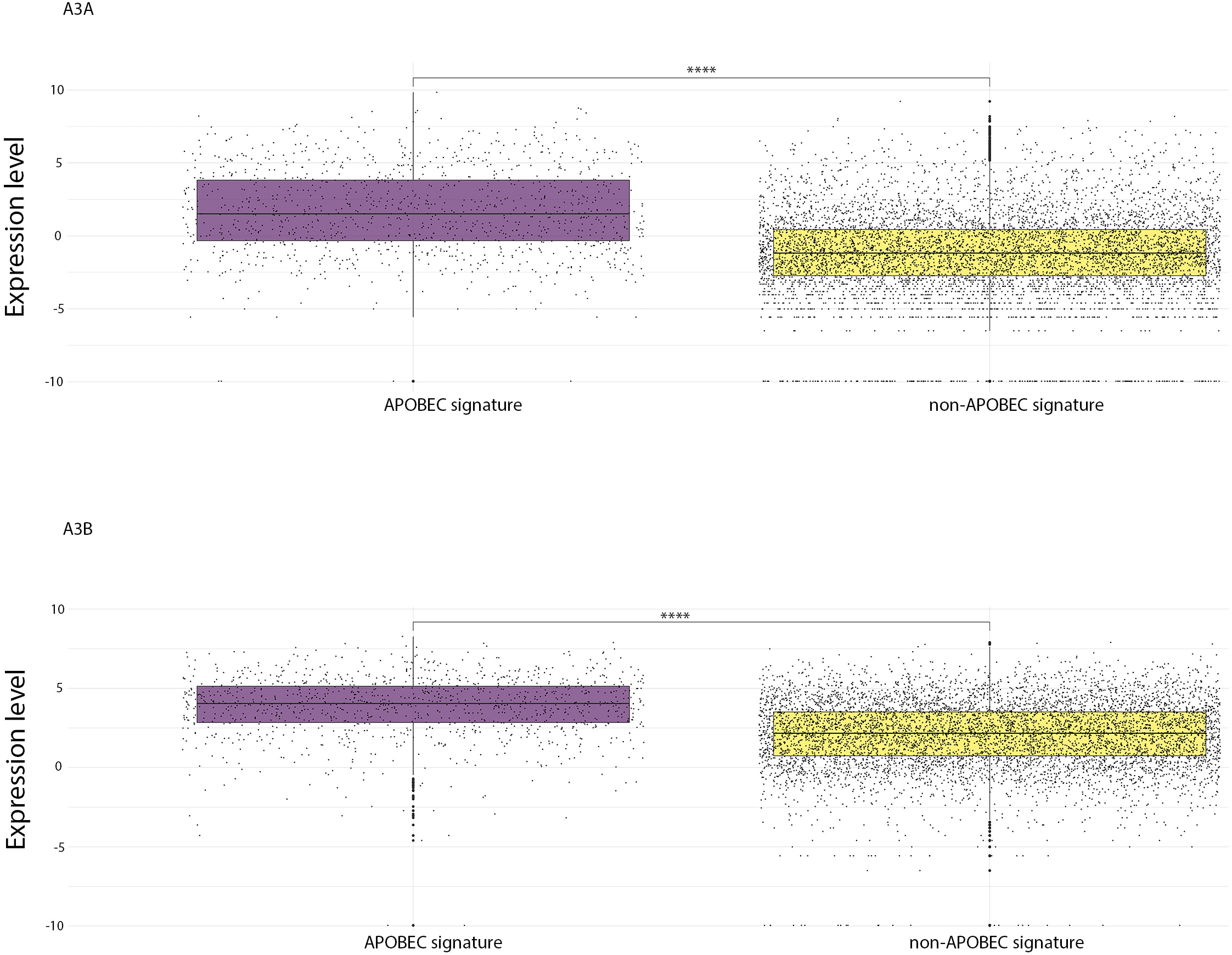
APOBEC3A and APOBEC3B expression level in patient samples with the APOBEC3-mutational signature. Gene expression of each APOBEC3 gene was compared in samples that do bear the APOBEC3 mutational signature vs. tumors that have non-APOBEC mutational footprints.

